# Deep Learning Initialized Compressed Sensing (Deli-CS) in Volumetric Spatio-Temporal Subspace Reconstruction

**DOI:** 10.1101/2023.03.28.534431

**Authors:** Siddharth S. Iyer, S. Sophie Schauman, Christopher M. Sandino, Mahmut Yurt, Xiaozhi Cao, Congyu Liao, Natthanan Ruengchaijatuporn, Itthi Chatnuntawech, Elizabeth Tong, Kawin Setsompop

## Abstract

**Introduction:** Spatio-temporal MRI methods enable whole-brain multi-parametric mapping at ultra-fast acquisition times through efficient k-space encoding, but can have very long reconstruction times, which limit their integration into clinical practice. Deep learning (DL) is a promising approach to accelerate reconstruction, but can be computationally intensive to train and deploy due to the large dimensionality of spatio-temporal MRI. DL methods also need large training data sets and can produce results that don’t match the acquired data if data consistency is not enforced. The aim of this project is to reduce reconstruction time using DL whilst simultaneously limiting the risk of deep learning induced hallucinations, all with modest hardware requirements.

**Methods:** Deep Learning Initialized Compressed Sensing (Deli-CS) is proposed to reduce the reconstruction time of iterative reconstructions by “kick-starting” the iterative reconstruction with a DL generated starting point. The proposed framework is applied to volumetric multi-axis spiral projection MRF that achieves whole-brain T1 and T2 mapping at 1-mm isotropic resolution for a 2-minute acquisition. First, the traditional reconstruction is optimized from over two hours to less than 40 minutes while using more than 90% less RAM and only 4.7 GB GPU memory, by using a memory-efficient GPU implementation. The Deli-CS framework is then implemented and evaluated against the above reconstruction.

**Results:** Deli-CS achieves comparable reconstruction quality with 50% fewer iterations bringing the full reconstruction time to 20 minutes.

**Conclusion:** Deli-CS reduces the reconstruction time of subspace reconstruction of volumetric spatio-temporal acquisitions by providing a warm start to the iterative reconstruction algorithm.

## 1 INTRODUCTION

Highly undersampled, spatio-temporal MRI acquisition techniques have enabled whole-brain multi-parametric mapping in incredibly short exam times. These techniques leverage highly-efficient k-space encoding^1,2,3^, as well as temporal subspace reconstruction^4,5,6,7,8,9,10,11,12^ with carefully-chosen regularization to achieve high-quality reconstruction without detrimental noise and undersampling artifacts despite the high rates of acceleration. However, this generally comes at the cost of long reconstruction times, especially for for high isotropic resolution volumetric imaging cases, making such methods difficult to integrate into clinical practice despite the high acquisition efficiency.

For example, consider the “Tiny Golden Angle Shuffling Spiral Projection Imaging Magnetic Resonance Fingerprinting” (TGAS-SPI-MRF^12^) volumetric acquisition, which will be the target application of this work, that demonstrated the use of an optimized trajectory to achieve whole-brain multi-parametric mapping at 1mm isotropic resolution in 1-2 minutes of acquisition time using the locally low-rank (LLR)^10^ regularized reconstruction implemented in BART^13^. With the settings of BART used by Cao et al.^12^, the reconstruction takes around two hours and 10 minutes using up to 80 CPU threads with around 140 GB of peak resident memory usage. The large dimensionality of the problem hinders out-of-the-box GPU utilization. This computational performance is achieved after BART’s default optimizations to improve reconstruction speed, such as utilizing the theoretically optimal Fast Iterative Shrinkage-Thresholding Algorithm (FISTA)^14^ for solving the LLR-regularized optimization, as well as the combination of the “Toeplitz Point Spread Function”^15,16,17^ and the “spatio-temporal kernel”^10^ to reduce the per-iteration compute time at the cost of an even more memory intensive pre-calculation (using up to 400 GB of resident memory).

Concurrently with the development of fast spatiotemporal MRI acquisition, significant progress has been made in utilizing deep learning for image reconstruction for acquisitions with high rates of acceleration^18,19,20,21,22,23^. These methods commonly leverage an “unrolled” deep learning architecture, where the algorithm alternates between performing network inference and enforcing a physics-based Data-Consistency (DC) step akin to traditional (compressed sensing like) iterative methods for solving regularized linear inverse problems^24,25,26,27,28,29^. These works fall under the “physics-driven” classification proposed by^23^, and will be referred to as such in the sequel. The integration of the DC term into these unrolled physics-driven methods has enabled the robust application of deep learning based MRI reconstruction without access to the copious amounts of training data typically required by other deep learning methods^18^. These methods have achieved excellent reconstruction performance at significantly faster processing times compared to their iterative convex algorithm counter-parts, which makes such frameworks a promising means of achieving fast spatio-temporal MRI reconstruction.

However, utilizing such unrolled physics-driven methods out of the box can be challenging depending on the dimensionality of the problem of interest. To use the target TGAS-SPI-MRF application as an example, the underlying desired signal is of dimensions 256 × 256 × 256 × 5, where 256 × 256 × 256 is the matrix size of the acquisition, and 5 denotes the number of subspace coefficients (that are needed to represent the temporal characteristics of the underlying signal)^4,6,8,10,11,12^. This, along with using multiple receive channels for the acquired data across k-t-space dramatically increases the dimensionality of the reconstruction problem, making such unrolled physics-driven reconstructions extremely computationally challenging to both train and deploy. This problem is also not separable in the 3D spatial dimensions, as the k-space trajectory is designed to spread aliasing along all spatial dimensions, unlike acquisitions with rectilinear trajectories and undersampling, where the problem can be divided along the readout dimension.

To give a concrete example, the current infrastructure at Stanford Health Care and the Lucile Packard Children’s Hospital (Stanford, CA, USA) have dedicated MRI reconstruction servers with multiple GPUs with memory capacities in the range from 6 GBs to 12 GBs. Preliminary exploration of leveraging multiple GPUs in parallel for image reconstruction were futile due to the synchronization costs (likely due to the large problem size of the application), and the 12 GB GPUs by themselves were not sufficiently large enough to utilize the unrolled physics-driven methods for the target application when utilizing a Residual Network^30^ with 3D convolutions to perform model inference.

With these constraints in mind, this work proposes Deli-CS, which stands for “Deep Learning Initialized Compressed Sensing”. The goal of this framework is the rapid and highly compute-efficient subspace reconstruction of spatio-temporal MRI acquisitions (such as TGAS-SPI-MRF) with the goal of clinical deployment. This framework thus targets less than 6 GB peak GPU memory usage and a reconstruction of the 2-minute TGAS-SPI-MRF acquisition that is comparable in quality to iterative LLR reconstruction of the same data. Unlike other DL methods that have been developed for de-noising or other ways of improving image quality, the aim of this project was strictly decreased reconstruction time and hardware constraints, as good image quality was already demonstrated for this application by Cao et al.^12^

By being GPU efficient, the proposed framework is expected to scale well to ultra-high resolution sub-millimeter applications such as the 0.66mm isotropic resolution “ViSTa-MRF” acquisition for high-fidelity whole-brain myelin-water fraction (MWF) and *T*_1_, *T*_2_ and proton density mapping on a clinical 3T scanner^31^. Additionally, enforcing a minimal amount of GPU memory for training simplifies the process of “continuous training” to update the learned model to account for potential distribution shifts, which is an important factor to consider when deploying deep learning methods^32,33,34,35,36^.

This manuscript is organized as follows: First, the traditional LLR regularized reconstruction is implemented in Python using SigPy^37^ in a GPU-efficient manner. Next, the Deli-CS framework is described, where a fast initial reconstruction (generated by multiplying the data with the adjoint of the forward operator, **A**, described below) is fed into a neural network that attempts to predict the final reconstruction. The training and inference are performed in a block-wise manner to significantly lower memory usage. Finally, since the model does not have an integrated DC term and is not unrolled (this would be classified as “data-driven” by Hammerik et al.^23^), data consistency is enforced by a “compressed sensing certification” step, which is hypothesized to converge faster with the DL initialization.

## 2 BACKGROUND

Spatio-temporal subspace-reconstruction^4,10,12^ is posed as a linear inverse problem where the acquisition operator, **A**, models the transformation from the input subspace coefficient images (**x**) to the data acquired (**b**). Let the subscript *t* denote the data acquired at the *t*^*th*^ TR, with *T* being the total number of TRs, and let *K* denote the number of coefficient images, or the rank of the low-rank subspace utilized. Then, the acquisition operator **A** is as follows:

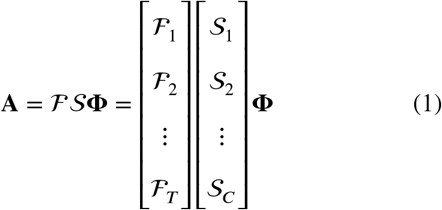

Here, ℱ denotes the forward NUFFT operator^38^, 𝒮 the projection onto coil sensitivity maps according to the SENSE^39,40^ model and Φ denotes the prior low-rank linear subspace with dimensions [*T* × *K*]. As shown above, the ℱ operator can also be expressed as a stack of smaller NUFFT’s, ℱ_*t*_, that each express the transform from image to k-space for the *t*^*th*^ TR. Similarly, 𝒮 can be expressed of a stack of *S*_*c*_ denoting the projection onto the *c*^*th*^ receive channel. This formulation will be used below when the memory efficiency optimization of the operator is discussed.

The low-rank subspace **Φ** is estimated by taking the Singular Value Decomposition (SVD) of signal dictionary generated from Bloch-simulations using the following realistic brain tissue parameters that match prior work^12^.

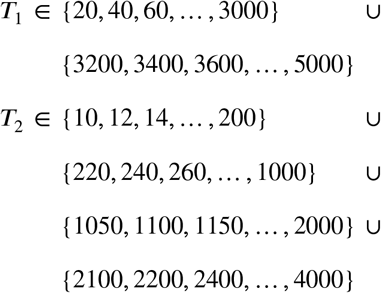

The forward operation Φ**x** recovers the TR images of the TGAS-SPI-MRF acquisition. A rank *K* = 5 subspace was deemed sufficient in capturing the signal variation as per Cao et al.^12^. Please see Cao et al.^12^ for more information about the acquisition and subspace forward model formulation.

The linear inverse problem used to solve the subspace reconstruction is as follows:

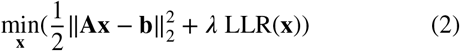

Here, *λ* is the regularization value and LLR denotes the locally low-rank constraint^10,12^.

## 3 METHODS

### 3.1 Data Acquisition

The method of TGAS-SPI-MRF proposed by Cao et al.^12^ was used, but a modification was made to the acquisition by replacing the RF excitation pulse with a water-exciting rectangular pulse with duration of 2.38ms to suppress the fat signal^41^.

In summary, the TGAS-SPI-MRF acquisition consists of an initial adiabatic inversion pulse followed by a 500 TR long readout train (TI/TE/TR = 20/0.7/12ms) with varying flip angles (10 to 75 degrees) and a rotating 3D center-out spiral trajectory. Each acquisition group takes approximately 7.5 seconds and 16 such readout trains are required for a 2-minute acquisition, whereas 48 are used for a 6-minute acquisition, which was treated as the gold standard in this work.

Additionally, prior to each TGAS-SPI-MRF acquisition, a 20 second, low resolution (6.9 mm isotropic) gradient echo (GRE) pre-scan with a large field-of-view (FOV) of 440mm was performed. This pre-scan was used for automatic detection of the head position within the large FOV, so that the TGAS-SPI-MRF data could be automatically shifted to ensure that the brain, as well as the top of the head and nose, was within the smaller FOV used for the TGAS-SPI-MRF reconstruction. The large FOV pre-scan was also used to pre-calculate a coil compression matrix, such that any signal originating from outside the TGAS-SPI-MRF FOV after shifting (e.g. shoulders) could be removed using region-optimized virtual coil compression^42^ (ROVir) estimated from the GRE with the interference region set to any area outside the target FOV for the TGAS-SPI-MRF acquisition.

### 3.2 Memory Optimization

This work uses the Python-based SigPy^37^ framework for the following experiments for its ease-of-use reconstruction whilst retaining control over GPU memory management. To reduce the memory requirements of **A**, the following standard optimizations are made.

First, the forward model **A** is split into smaller blocks so that each block can be iteratively and independently efficiently evaluated on the GPU. Second, to avoid expanding into the range space of Φ (*T* ≫ *K*), the commutativity of the subspace operator Φ and the NUFFT is used akin to the spatio-temporal kernel leveraged in *T*_2_−Shuffling^10^. These two optimizations together result in the following forward model:

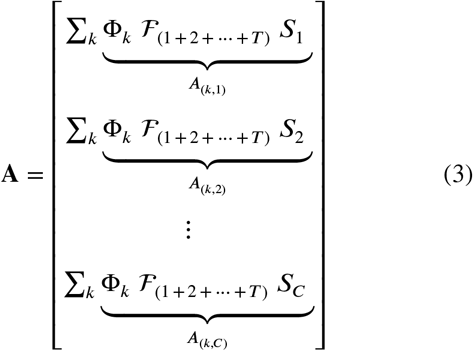

Here, Φ_*k*_ denotes the *k*^*th*^ column of Φ, and *S*_*c*_ denotes the *c*^*th*^ SENSE^39,40^ coil-sensitivity map. The individual smaller block linear operators, *A*_(*k,c*)_, can be evaluated one-by-one and thus reduces memory requirements. The input to each block linear operator, **x**_*k*_, is first transferred to GPU memory before applying the operator, and the resulting output, **A**_(*k,c*)_**x**_*k*_, is then transferred into CPU memory before the stacking over coils (*c*) and sum over coefficients (*k*) is applied.

The modified linear operator proposed in (3) achieves a computational speed of approximately 58 seconds per iteration of FISTA^14^ when solving (2) for a 2-minute TGAS-SPI-MRF acquisition in SigPy on a single GPU. The seconds-per-iteration of FISTA achieved by BART on the CPU is approximately 26 seconds. These times were observed on a Linux workstation with 80 threads on Intel Xeon Gold 5320 CPUs at 2.20GHz and an NVIDIA RTX A6000 GPU. Note that this performance is expected to vary between hardware.

However, the modified linear operator only utilizes approximately 13 GB of peak CPU memory and 4.7 GB of peak GPU memory, compared to 140 GB peak CPU memory for the BART implementation. BART uses Toeplitz embedding^16^, which reduces per-iteration computation time as the NUFFT’s in the acquisition model can be replaced by FFT’s, but increases memory usage and requires a step before iterations can start, which had a peak memory usage of >400 GB. Additionally, as methods are striving for ever higher resolution, the memory requirements for the BART reconstruction would grow well over what is available on most research computers, not to mention what is available in clinical settings.

### 3.3 Density Compensation

Having achieved considerably lower memory usage, the next optimization targets improving the iterative convergence of FISTA. The acquisition operator **A** is ill-conditioned in that the difference between the largest and smallest eigenvalue of **A**^∗^**A** is large, yielding slow iterative convergence. Assuming 25 seconds per iteration and 300 iterations, which was used by Cao et al.^12^, this results in approximately 2 hours 10 minutes required to reconstruct data. To improve this, first Pipe-Menon^43^ density compensation was integrated into the optimization (equation 2) as described in^17,44,45^, yielding the following optimization:

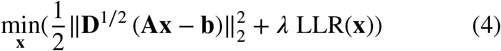

Here, **D** is the Density Compensation array designed to target ℱ_(1+2+…+*T*)_ in equation 3 so that **A**^∗^**DA** has better conditioning. In the block linear operator form, this becomes:

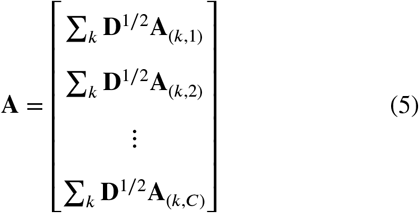

Similarly to the prior section, the sum across *k* and stacking across *c* is performed in CPU memory with each block evaluated in GPU memory.

While the inclusion of **D** does in principle cause noise coloring, in practice, careful tuning of the LLR regularization value was found to provide suitable levels of denoising, resulting in high quality reconstructions in fewer iterations. This reflects the results discussed in Baron et al.^17^, and demonstrates that Density Compensation can be leveraged to achieve high quality subspace reconstruction. The inclusion of density compensation into the optimization formulation is seen to significantly reduce the number of required iterations, with convergence reached after only 40 iterations (39 minutes).The reconstruction quality was also visually sharper compared to the 300 iterations (2 hours and 10 minutes) presented in Cao et al.^12^ In this work, the LLR block size used was 8.

### 3.4 Field of View Processing

It is beneficial to reduce the image size of the reconstruction for lower memory usage and faster processing times. However, using a smaller FOV is not advisable if there is signal originating outside the reconstructed FOV as it results in artifacts during the reconstruction. For the TGAS-SPI-MRF brain imaging application, signal from the shoulders and neck are particularly troublesome. This is overcome by utilizing the automatic FOV shifting approach proposed in Baron et al.^17^ that is augmented with a newly proposed coil compression method for additional robustness^42^.

The automatic FOV centering was done by reconstructing the fully sampled, Cartesian, GRE image using a Fourier Transform, and performing a sum-of-squares combination of data from the multiple receive coils. The image was then flattened by taking a maximum intensity projection through the sagittal plane. The resulting 2D image was then smoothed, binarized, and a bounding box was calculated around the largest continuous area using the OpenCV toolbox^46^. A shift was then calculated to ensure the top and front of the bounding box was within the target field of view.

Baron et al.^17^ derived the binary image from the eigenvalues of the estimated ESPIRiT^40^ maps, which is not used in this work as the target TGAS-SPI-MRF application utilizes 48 receive coils during the acquisition, making ESPIRiT computationally expensive. Additionally, the FOV shifting is performed *before* applying any coil compression techniques, which enables the shifted GRE image to be used as a reference for further suppression of signals originating outside the FOV using the ROVir method^42^ as outlined below.

The data was pre-whitened as described by Kim et al.^42^ before ROVir was applied. Signal outside the field of view was removed by throwing away 8 virtual ROVir coils that contained the most signal in the interference region. After that, SVD compression to 10 virtual channels of the remaining 40 channels were performed to reduce the size of the computation. The complete coil compression matrix containing coil whitening, ROVir, and SVD compression was calculated based on the GRE, and used for compression of the TGAS-SPI-MRF data.

The reconstruction, automatic FOV shifting, and coil processing matrix calculation for the GRE data takes less than 30 seconds, and can be run while the TGAS-SPI-MRF data is being acquired.

### 3.5 Deli-CS

With Deli-CS, the goal is to achieve reconstruction of TGAS-SPI-MRF data that is of comparable quality to the traditional reconstruction with minimal memory footprint as well as faster convergence than previously demonstrated. To achieve this, the following design pillars are utilized:

1. *Fully Leveraging MRI Physics*. The aim is to solve the traditional model based reconstruction and only using DL as a way to “kickstart” the reconstruction. As a first step, an approximate reconstruction is performed by gridding the acquired data using the adjoint of the forward operator (**A**^∗^**b**). By not using an iterative reconstruction (e.g. conjugate gradient) and not regularizing, the input to the next step suffers from severe temporal-aliasing artifacts and increased noise, in particular in the lower energy subspace components, which the following deep learning step is expected to robustly mitigate. The resulting subspace coefficient images will be referred to as “Deli-CS Input”.
2. *Block-based Data-Driven Deep Learning*. A deep learning network is trained to denoise the input image. The model is both trained and deployed in a “block-wise” manner to reduce the memory and training data requirements, and is consequently *not* integrated with DC terms in an unrolled manner. This also avoids back-propagation through the high-dimensional acquisition operator **A** during training, which is challenging to do even with a GPU with large memory capacity. The block-based processing proposed in this step allows the deep learning model to be trained and deployed efficiently with 5 GB of GPU memory. The resulting inference will be referred to as “Deli-CS Prediction”.
3. *Compressed Sensing Certification*. Since the above network is block-based and data-driven, the inferred reconstruction is susceptible to hallucinations. The inferred result is therefore only used to initialize equation 4 solved with an iterative reconstruction. By initializing the iterative reconstruction with the inferred result, the number of iterations required to converge is significantly reduced. Additionally, the resulting image is still “Compressed Sensing Certified” in the sense that the resulting image satisfies the same convergence criterion as the traditional reconstruction achieved when solving equation 4. This step will be referred to as the “refinement” step, with the resulting reconstruction denoted “Deli-CS Refined”.

The full Deli-CS pipeline is depicted in figure 1.

**FIGURE 1.**
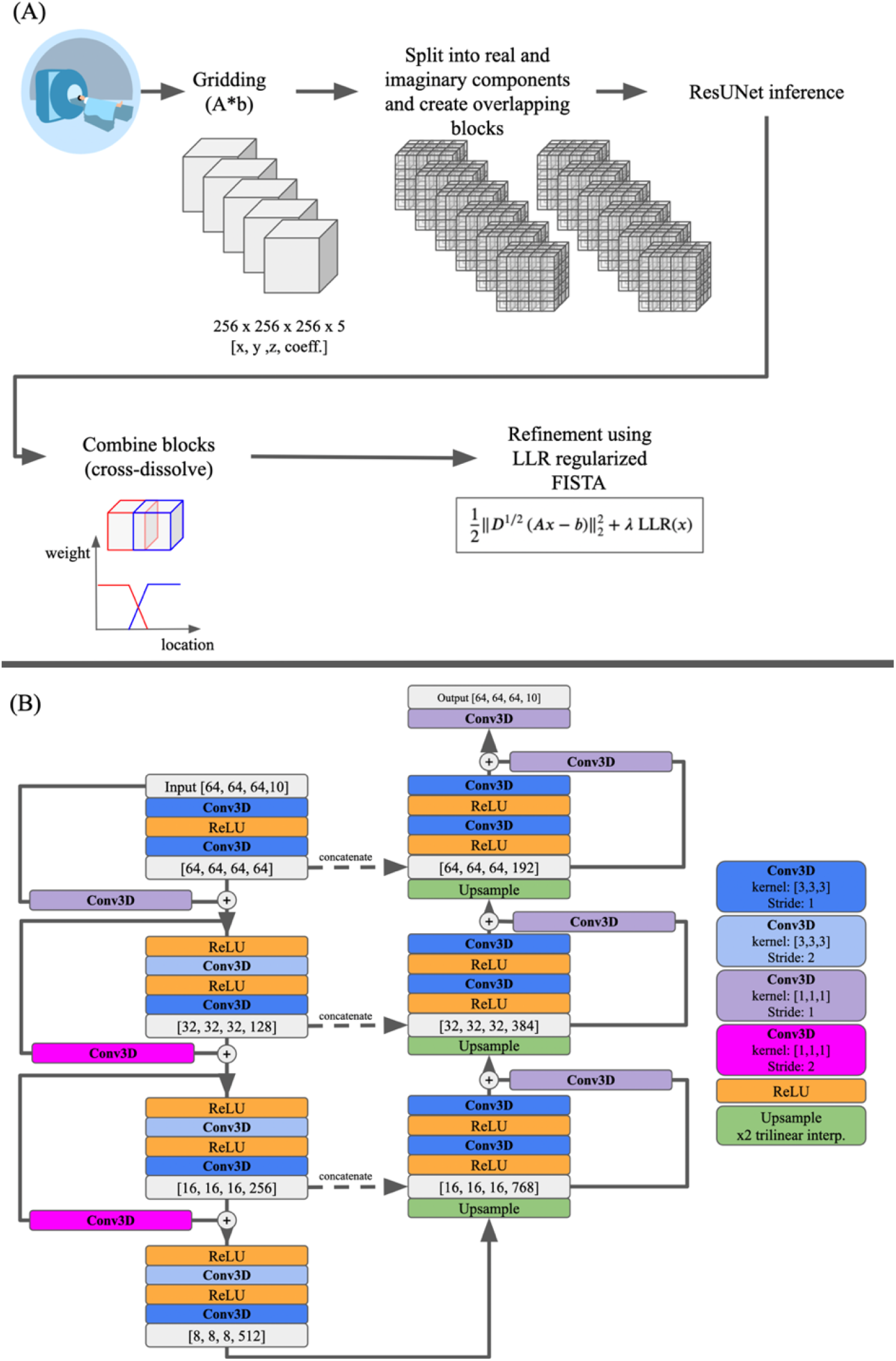
The Deli-CS pipeline is depicted in (A) with further detail regarding the ResUNet shown in (B). The gray boxes in (B) show the data size at various layers of the network and the first three numbers are the spatial dimensions, whereas the fourth number is the number of feature channels.

### 3.6 Basis Balancing

The subspace basis is estimated by taking the SVD of a dictionary of realistic signal evaluations generated using the Bloch equation and using the singular vectors corresponding to the top five singular values to form Φ^12^. This rank-truncation level was deemed sufficient in capturing the signal variation as per Cao et al.^12^, and the parameters used to derive the dictionary was presented in section 2. Using this basis directly results in very low signal level in the fourth and fifth coefficients as most of the signal is already captured in the first three basis components. This is shown in figure 2(A) and (B). To ensure each coefficient image contributes roughly equally to the objective function when model training and to more equally distribute the artifacts across coefficients, a “basis balancing” heuristic is proposed to equally distribute the energy across all the coefficients.

**FIGURE 2.**
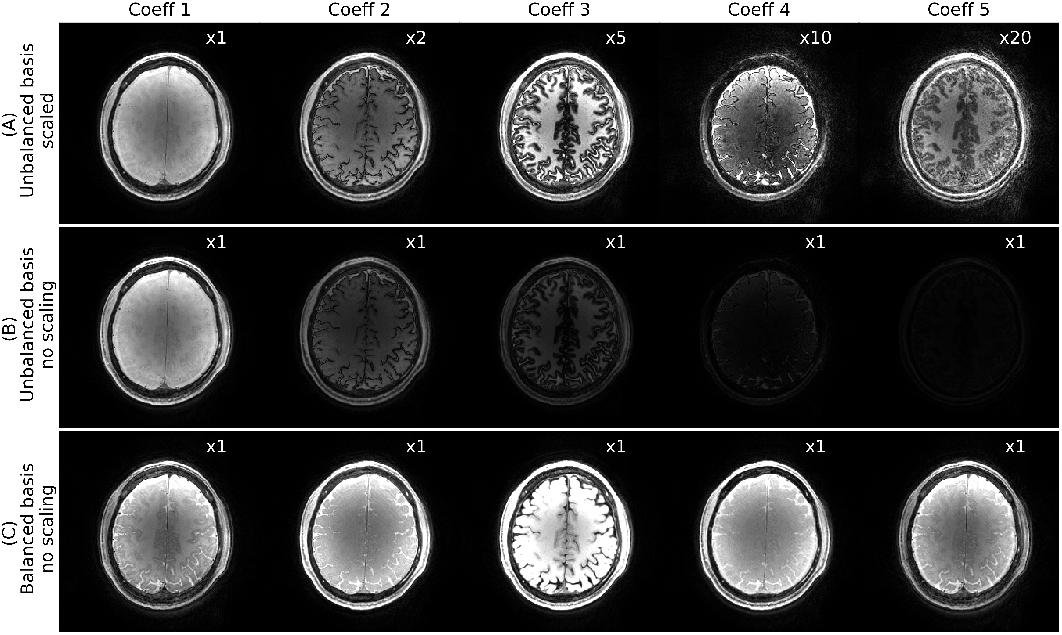
This figure depicts how the 6-minute gold standard images (magnitude only) look without basis balancing (A,B), and with basis balancing (C). Note variation in image intensity between bases in (A,B) which could cause instabilities in the subsequent DL step. (C) looks like it has less variability between bases, but the balancing method transfers tissue information to the image phase, which is not depicted here.

The low-rank basis Φ of dimensions *T* × *K* with Φ_*k*_ denoting the *k*^*th*^ column of Φ. Let *B* denote the new basis with *b*_*k*_ denoting the *k*^*th*^ column. In order to balance *B*, the columns *b*_*k*_ are derived so that *b*_*k*_ consists of equal contributions from each column of Φ. This translates to an equality constraint on the magnitude of the inner products. That is to say, for some constant *a*,

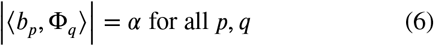

In other words, *B*^∗^Φ is a matrix where the magnitude of each matrix is the same.

One operator that satisfies this property is the Discrete Fourier Transform (DFT) matrix. Let Θ be the DFT matrix of dimensions *K* × *K*. Then, the balanced basis *B* can be derived as:

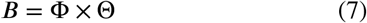

With Θ normalized to have unitary columns, *B* is also an orthonormal matrix. Additionally, since the columns *B* are constructed from a linear combination of the columns of Φ, *B* and Φ span the same subspace. In other words, *B* is a linear combination of the columns of Φ and thus a reconstruction with or without basis balancing should give the same result up-to a change-of-basis transformation. Since *B* is derived from Θ, Θ^−1^ can be used to perform a linear change of basis from *B* to Φ.

All reconstructions presented here used the balanced basis *B*, but the coefficient images with respect to Φ are shown after applying the change-of-basis operation Θ^−1^ to the recovered coefficients. This is done for ease-of-comparison against prior reported reconstructions such as in Cao et al.^12^ Using *B* for the initial reconstruction yields coefficient images of roughly equal signal level with no one coefficient suffering from significantly more artifacts compared to the others as shown in 2(B).

### 3.7 Experiments

To train and verify the Deli-CS framework, data from 14 healthy volunteers were acquired on a 3T Premier MRI scanner (GE Healthcare, Waukesha, WI) and a 48-channel head receiver-coil. GRE and TGAS-SPI-MRF data were acquired. The TGAS-SPI-MRF acquisition time was 6 minutes, acquired resolution was 1 mm isotropic, and FOV was 220 mm isotropic. The data was retrospectively sub-sampled to simulate a 2-minute acquisition. The data were partitioned as 10 training, 2 validation and 2 testing subjects. The technique was additionally tested on three sets of patient data (all male, ages 28, 49, and 75) acquired as additional scans to the standard of care protocols at a local outpatient imaging center using two different 3T Signa Premier scanners. The patients were scanned with a prospectively accelerated 2-minute TGAS-SPI-MRF protocol and thus no 6-minute reference comparison was possible for these cases.

All human data were acquired with informed consent using protocols approved by the local institutional review board at Stanford University.

Coil sensitivity maps were estimated with JSENSE^47^ from the time averaged acquisitions to maintain fast reconstruction time (JSENSE coil sensitivity estimation took approximately 30 seconds compared to over 10 minutes using ESPIRiT). The dictionary and subspace were generated as described in section 2 above, and finally template matching was used to estimate the (*T*_1_, *T*_2_) parametric maps^1^.

The reported *λ* values for all reconstructions are after right-hand-side of the DC term of equation 4 (i.e. *D*^1/2^*b*) is normalized to have unitary *l*_2_−norm.

A gold standard LLR reconstruction was performed on the 6-minute data with an LLR block size of 8 and a *λ* value of 3 × 10^−5^ with 40 FISTA iterations. The matrix size was 256 × 256 × 256. This “gold standard” reconstruction was only used for reference to compare performance of the different reconstruction approaches of the retrospectively undersampled 2-minute case.

The retrospectively undersampled 2-minute data LLR reconstruction was also performed using an LLR block size of 8 and a *λ* value of 5 × 10^−5^ with 40 FISTA iterations. In both the 6-minute and 2-minute case this was sufficient for convergence. The assumed matrix size was 256 × 256 × 256. These data were used as target reconstructions when training the Deli-CS network.

For training the Deli-CS network, corresponding blocks of dimensions 64 × 64 × 64 × 5 were extracted from the initial adjoint reconstruction and the reference reconstruction of the 2-minute acquisition. For data augmentation, random flips, transposes, absolute scaling, and shifts were performed.

After global normalization, if the block’s DC coefficient’s standard deviation was below 0.3, the block was discarded. This simple filter ensures that the network avoids learning from regions with no signal variation (e.g. background only blocks). The blocks are subsequently split into real and imaginary components, and concatenated along the subspace dimension. A total of 623 training blocks from 10 subjects, and 152 validation blocks from 2 subjects were used. These blocks are piped into a ResUNet^48^ with 3D convolutions, where the channel dimension corresponds to the subspace dimension. The ResUNet utilized 3 residual encoding blocks and 3 residual decoding blocks with a filter size of 3 and ReLU activation. The total number of trainable parameters was 23.8 million. The model was implemented in PyTorch Lightning^49,50^ and trained for 479 epochs (stopping criteria defined by minimal validation loss) using the Adam optimizer^51^ with a learning rate of 1 × 10^−5^. The training utilized 5 GB of GPU memory.

During inference, blocks of spatial dimensions 64×64×64 were extracted from the initial reconstruction with a 16 voxel overlap in all dimensions. The resulting blocks were combined by applying a linear cross-blend operation to the overlap region to smooth out inconsistencies between blocks at the block edges.

For the final step of Deli-CS, the model prediction is used to initialize equation 4. The iterative reconstruction used for refinement utilized the same parameters as the reference 2-minute reconstruction.

Image quality in both the estimated coefficient images and the resulting quantitative maps was initially assessed qualitatively. Secondly, the T1 and T2 values for different number of reconstruction iterations were assessed in a gray matter and white matter mask generated by FSL FAST^52^ after brain segmentation using FSL BET^53^.

## 4 RESULTS

Using Density Compensation and GPU optimized processing improves both reconstruction quality (sharpness in particular) and time compared to the prior work that used LLR^12^ without density compensation. This is shown in figure 3.

**FIGURE 3.**
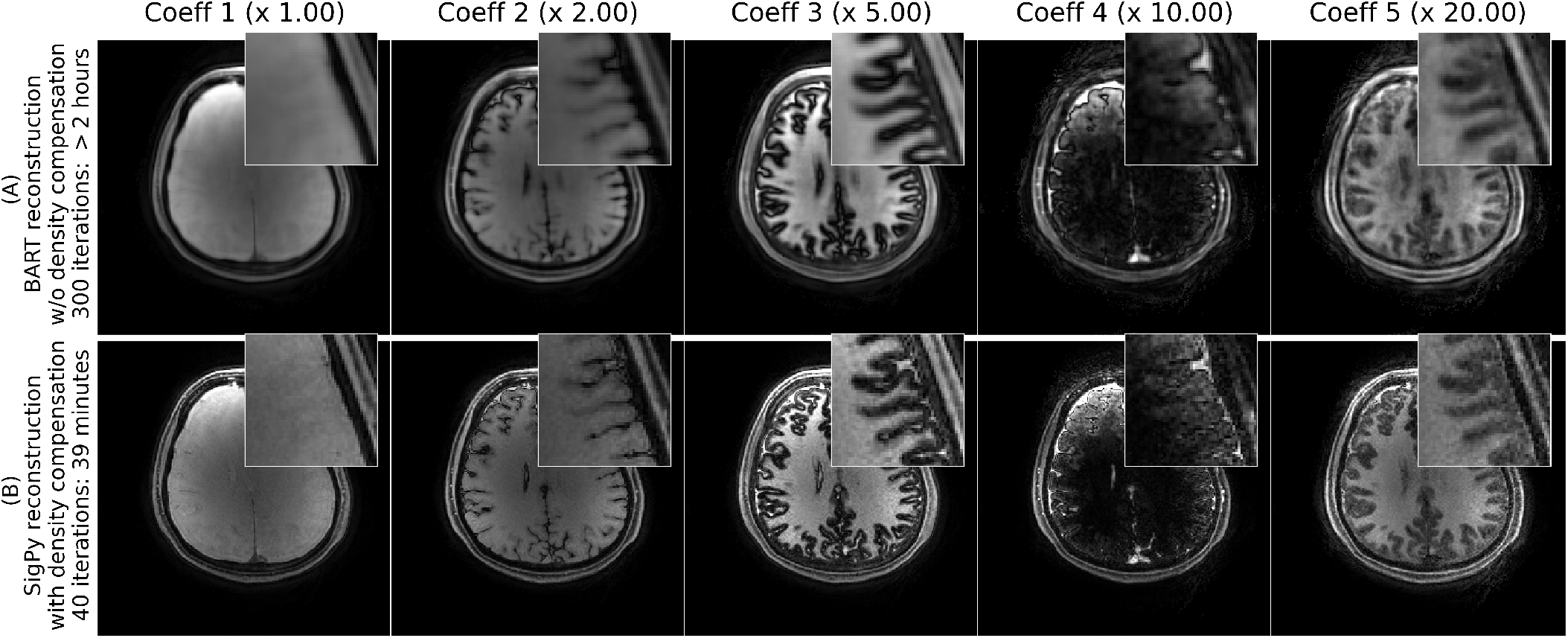
This figure compares a prior method by Cao et al.^12^ (A) to the updated reconstruction (B) that includes density compensation to significantly improve the conditioning of the problem, resulting in both much faster iterative convergence and improved sharpness compared to not using density compensation. The data shown here is for a 2-minute acquisition.

As shown in figure 4(A) and (B), the Deli-CS initialized prediction of T1 maps had a slightly lower RMSE error than the reference 2-minute LLR reconstruction after 4 iterations and after 20-30 iterations both reconstructions reached convergence. However, for the Deli-CS initialized reconstruction, the iterations mostly removed local biases, whereas the conventional iterative reconstruction mostly denoised the T1 maps. T2 maps estimation on the other hand benefited greatly from the Deli-CS initialization as can be seen in figure 4(C) and (D). The convergence of the T2 values for the conventional reconstruction required the full 40 iterations, whereas the Deli-CS initialized estimates had low error from the beginning and the iterative refinement changed the error from bias to more noise-like error.

**FIGURE 4.**
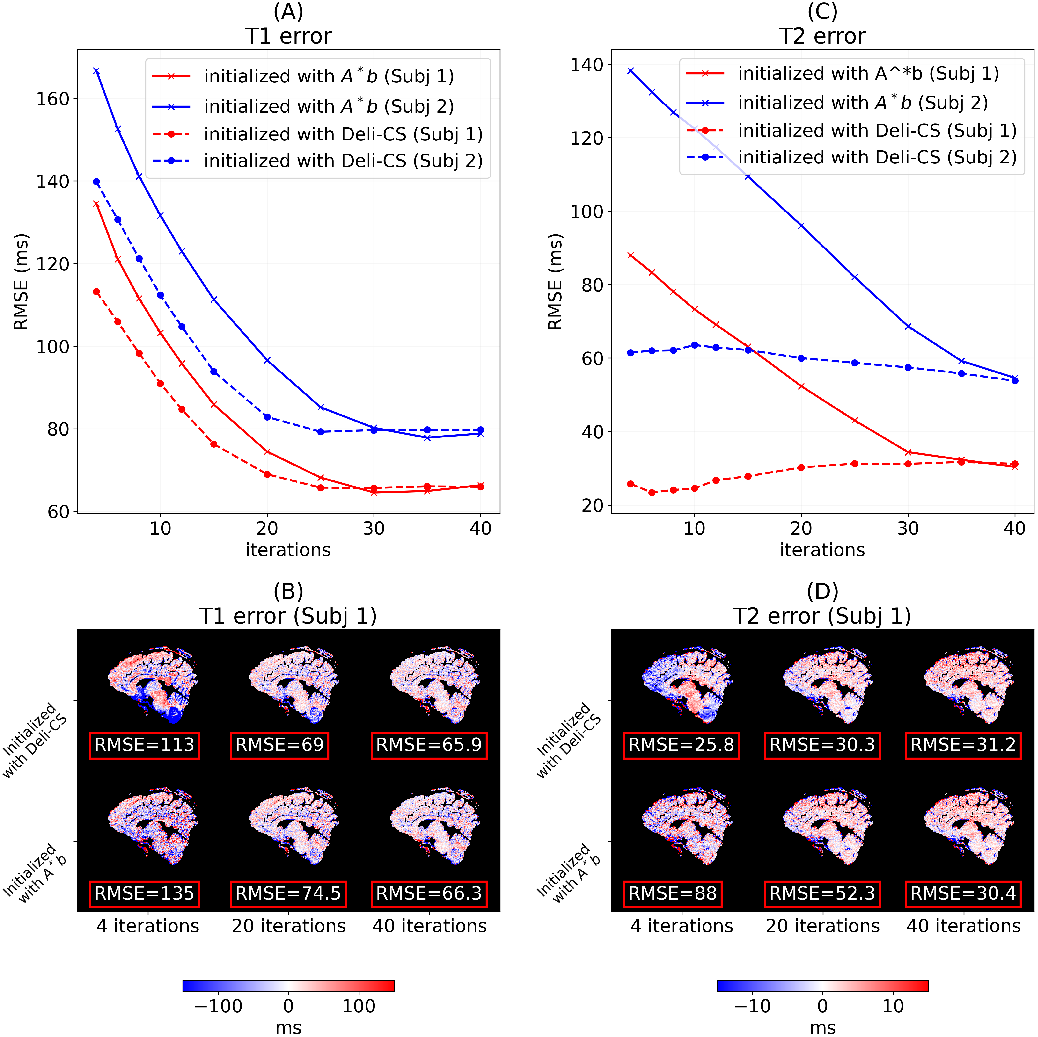
The estimated T1 RMSE values (A,B) and especially T2 RMSE values (C,D) within the white matter and gray matter in the two test subjects are shown to converge faster for the Deli-CS approach than standard initialization of the FISTA reconstruction.

After 20 iterations of refinement, the RMSE error of the T1 and T2 maps were similar to that of 40 iterations of adjoint-initialized reconstruction, providing a 2x acceleration factor on top of what the memory efficient and density compensated SigPy implementation of the algorithm already generated.

The reference iterative reconstruction took 39 minutes; the Deli-CS initial adjoint reconstruction took approximately 20 seconds; the Deli-CS model inference took approximately 20 seconds; the final refinement step with 20 iterations took approximately 19 minutes and 20 seconds.

Thus, the initialization approach enables approximately > 2*x* faster processing times with 50% fewer iterations.

Figure 5 shows a sagittal view of all steps within the reconstruction pipeline as well as the T1 and T2 quantification for one test subject. A magnitude threshold was applied to the figures to zero-out regions outside the FOV of the signal for presentability. The DL step smoothed the image significantly, which can be seen more clearly in figures 6 and 7, and caused some bias in the quantitative values especially in low signal areas around the brain stem (figure 5). The refinement step retrieved more sharpness and correct quantitative values in the low signal areas.

**FIGURE 5.**
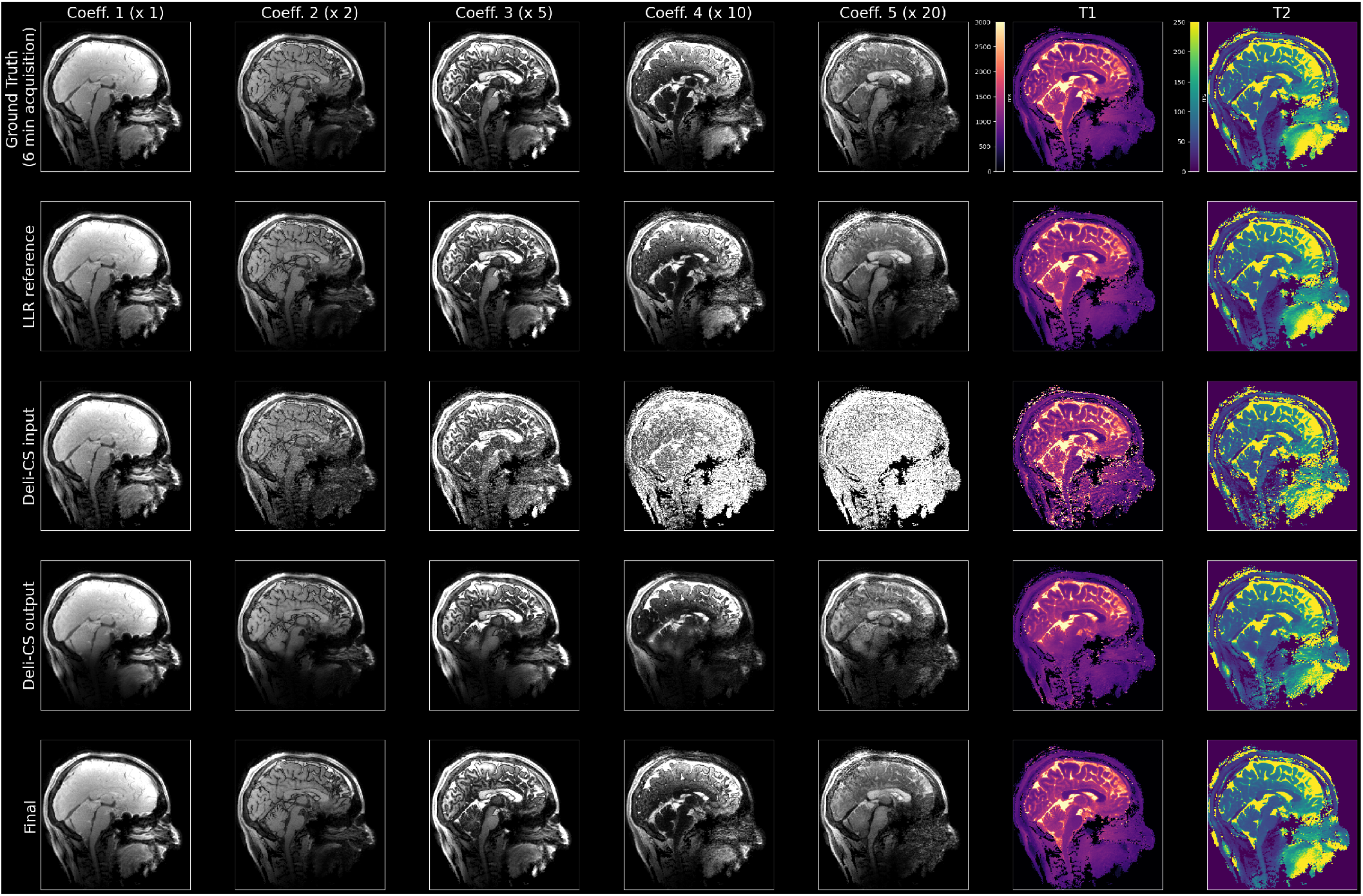
This figure compares the coefficient sagittal images recovered from the TGAS-SPI-MRF data using various methods for one of the healthy test subject. The first row denotes the reference LLR reconstruction of the 6-minute data acquisition, and the second row denotes the LLR reconstruction of the retrospectively under-sampled 2-minute acquisition. The remaining rows depicts the various steps of the Deli-CS pipeline. The third row shows the initial adjoint reconstruction, the fourth row shows the model inference, and the fifth row shows the reconstruction after iterative refinement.

**FIGURE 6.**
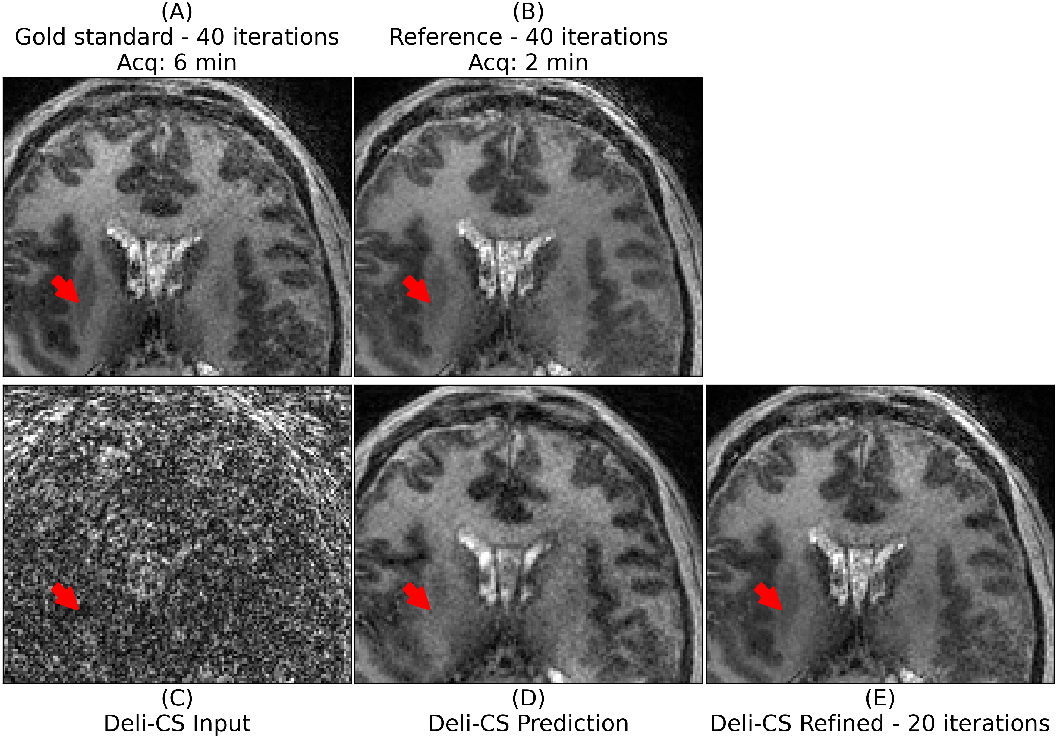
In this axial view of test subject 2 of the fifth (lowest energy) coefficient using the un-balanced basis that contains much of the white and gray matter contrast of the contrast of the putamen (denoted with a red arrow) is somewhat reduced using the Deli-CS inference but restored to the same level of contrast as the 2-minute acquisition by the refinement step. It is also noticeable just how much denoising the network performs.

**FIGURE 7.**
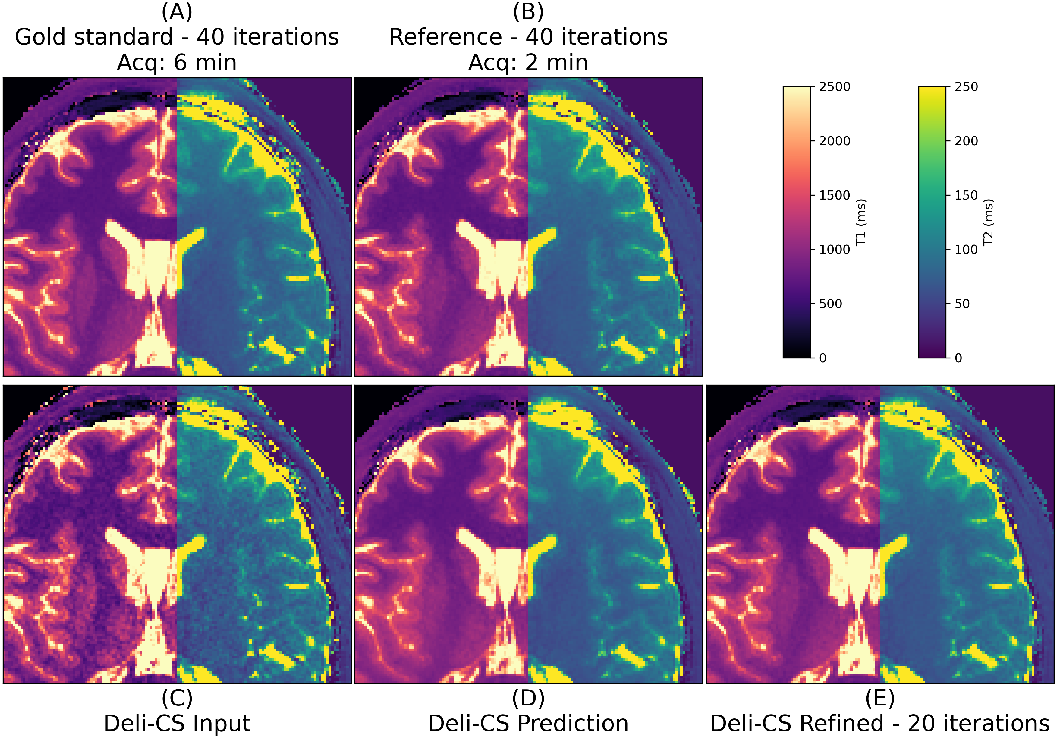
The quantitative maps of test subject 2 show high sharpness and contrast in both T1 and T2 maps with half the number of iterations when initialized with Deli-CS. T2 estimates benefit more than T1 estimates from Deli-CS initialization as they depend more on the lower energy coefficients.

As evidenced by figures 6 and 7, the refinement step adds missing features that were over smoothed or misassigned from the network prediction, mostly in the T1 map.

Similarly, in the patient data (figures 8, 9, and 10), great improvements in denoising by the network and sharpness and contrast from the refinement step was observed. The baseline quality of the 2-minute reference reconstruction was, however, lower than that of healthy volunteers. This is likely due to higher levels of motion as well as positioning further out of the head coil for patient comfort.

**FIGURE 8.**
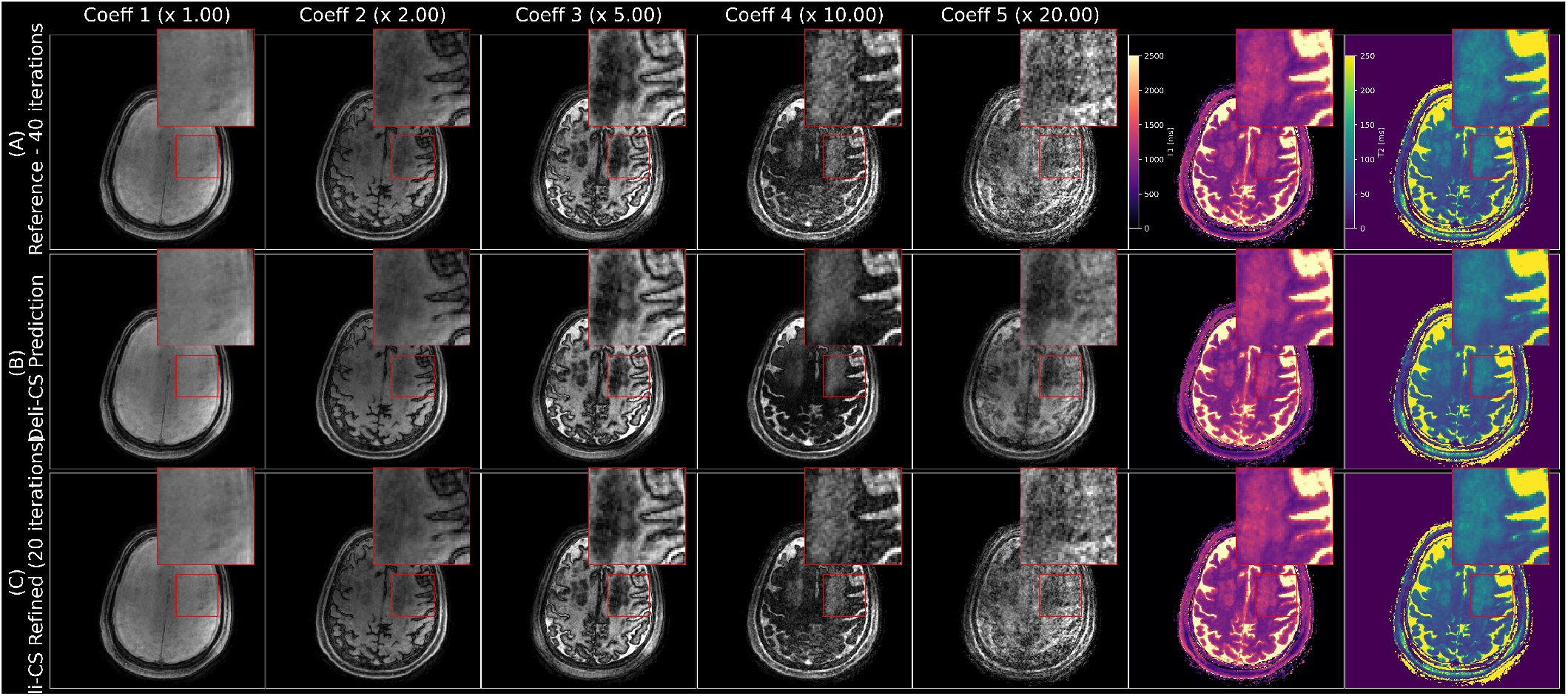
In this axial view of the subspace coefficients of a 75 year old male patient with chronic small vessel disease that affects both T1 and T2 in the deep white matter of the bilateral centrum semiovale (left side shown in the inset) is clear. Even though the original data in the lower energy coefficients is very noisy, a clean estimate is generated by the Deli-CS prediction step and the refinement step removes any bias generated by the Deli-CS prediction and confirms consistency with the acquired data. More detailed view available in figure 9.

**FIGURE 9.**
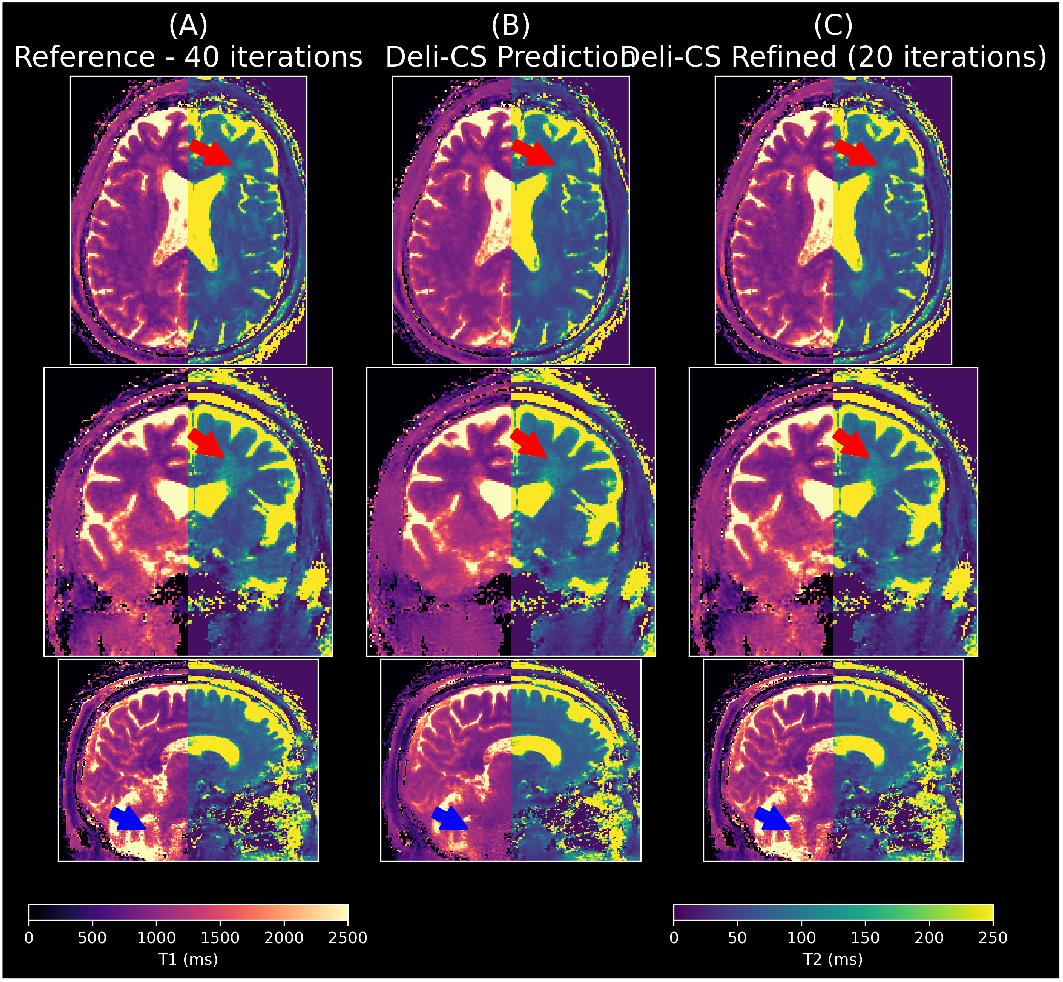
The refined quantitative maps in this patient (75 y/o male with chronic small vessel disease) show high sharpness and contrast in both T1 and T2 maps. Some areas are biased and blurred after the network inference (cerebellum: blue arrows, white matter lesion: red arrows), but recovered by the iterative refinement.

**FIGURE 10.**
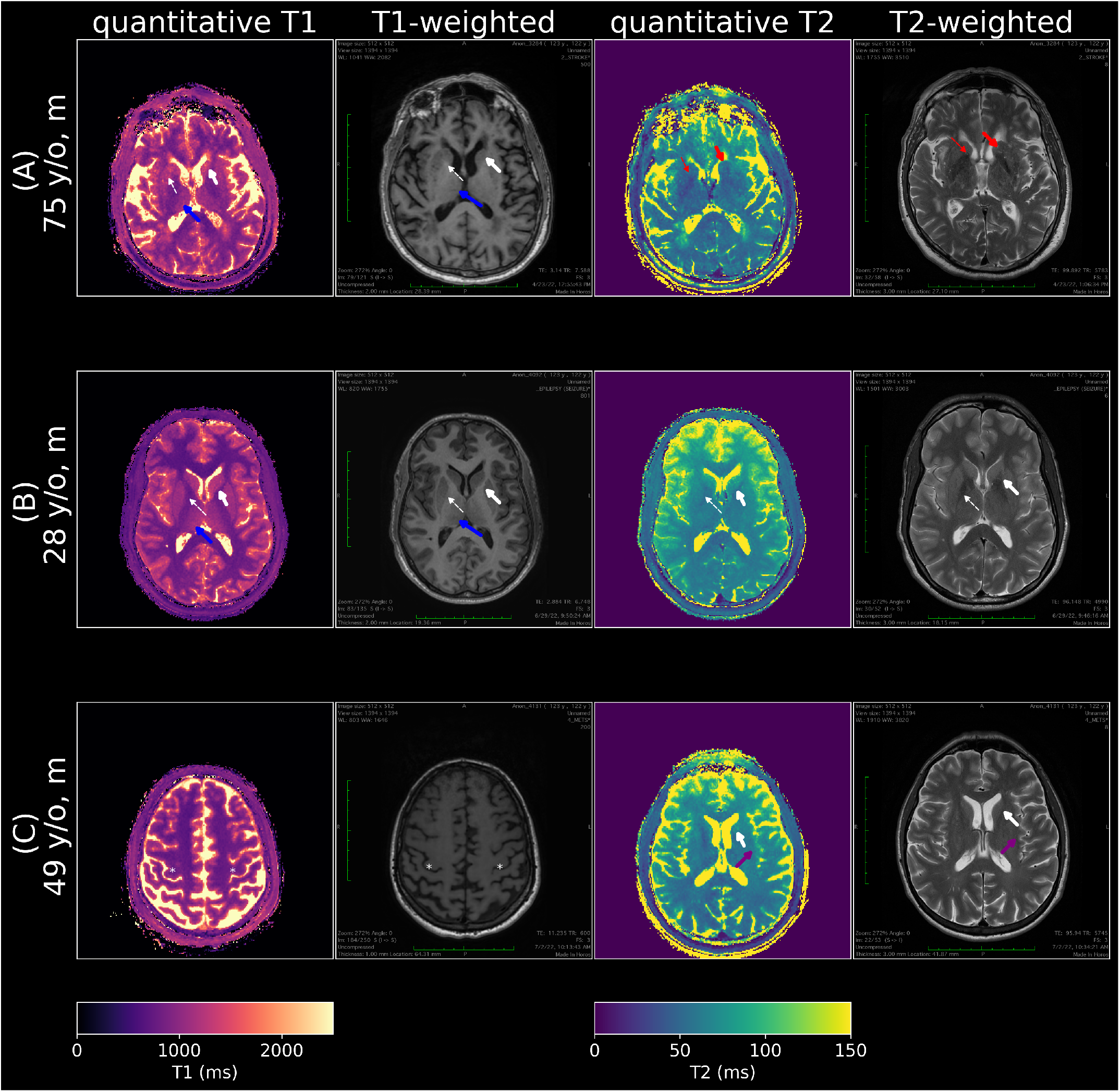
Comparison with standard clinical scans (slices approximately aligned). Note that T1 weighted images have inverted contrast to T1 maps as long T1 leads to low signal intensity on a T1 weighted image. In the T1 maps and T1 weighted images of (A) and (B) the caudate head (white arrow) and putamen (white dashed arrow) can be seen on the quantitative T1 map, but the delineation is slightly obscured compared to the T1 weighted image. The bilateral thalami (blue arrows) are more conspicuously delineated on the quantitative T1 map. The quantitative T1 map in (C) also demonstrates good gray-white distinction in the cerebrum as well as delineation of the hand-knob region (white asterisks). In (A) The striatum (red arrow) and globus pallidus (red dashed arrow) is somewhat obscured on the quantitative T2 map compared to the T2 weighted image. In (B), the quantitative T2 map distinguishes the caudate head and putamen well and the borders of the ventricles are well defined. In (C) the quantitative T2 map demonstrates clear borders of the ventricles. However the definition of the caudate head and insula (purple arrow) are somewhat obscured.

The 75 year old patient had indications for chronic small vessel disease in both the T1 and T2 maps shown in figures 8, and 9, whereas the other two patients exhibited no significant clinical findings, consistent with their standard-of-care imaging protocol as shown in figure 10.

The reference images in the clinical case are of somewhat lower quality than the healthy volunteers. This is likely due to motion and imperfect positioning in the head coil (further from receive coils and isocenter). However, despite not having trained on any data containing pathology, the Deli-CS network retained many clinically relevant features as shown in figure 10. Some deep gray matter areas that were visible on T2 weighted images were, however, obscured on quantitative T2 maps. This could be because the T2 weighted images have additional contrast from T1 and/or MT effects that are not captured in pure T2 maps^54^.

## 5 DISCUSSION

This work presented a framework for MRI reconstruction that targets high dimensional applications like volumetric non-Cartesian spatio-temporal subspace reconstruction, with the goal of reconstructing said application in clinically feasible time frame with modest hardware requirements. This was achieved with a block-based deep learning initialization approach, where the deep learning prediction was used to jump-start a regularized linear inverse problem. In clinical workflows this could allow the technologist/radiographer that acquires the scan to get the initial reconstruction or Deli-CS prediction to make an initial assessment on whether the scan was successful or need to be reacquired. The refined reconstruction, which takes an additional 20 minutes to reconstruct can then be sent to the radiologist for more detailed assessment.

The new, GPU optimized and sampling density compensated reconstruction method outperforms the prior reconstruction approach by Cao et al.^12^ as evidences by figure 3. The Deli-CS inference step efficiently denoises and smooths the initial reconstuction, and the refinement step suppresses hallucinations and bias, as well as over-smoothing. In principle, the optimization in 4 has a unique solution that the iterations will converge to, regardless of initialization. We noticed that the iterations when initializing with **A**^∗^**b** mainly denoises the result, whereas initializing with Deli-CS reduces bias in the quantitative parameter maps. The T2 maps benefit most from Deli-CS initialization as the T2 estimation is more dependent on the lower energy coefficient maps which suffer the most from noise and aliasing in the **A**^∗^**b** initialization.

In the patient data the denoising effect of the DL prediction step was very strong and the Deli-CS predictions before the refinement looked much cleaner than both the reference image and subsequent refinement step. But although the reconstruction was clean there is no way to certify whether or not the DL method hallucinated features that were not actually there and not even consistent with the data acquired as no data consistency is included in this step. As unrolled methods get more powerful and multi-GPU solutions become more readily available, the Deli-CS prediction can be used as a prior instead of a starting point in iterative reconstructions and thus be useful in improving image quality as well as reconstruction times.

The good recovery of the fourth and fifth coefficients in the healthy volunteer data is also expected to improve more advanced quantitative parameter fitting, such as in multicomponent modeling and multidimensional correlation spectroscopic imaging^55^. In particular, since a voxel typically consists of multiple tissue types, the better resolved fourth and fifth coefficients are expected to allow for the better fitting of multiple *T*_1_ and *T*_2_ values per voxel.

While this work aims to describe a general deep learning initialization approach, prior work^56^ has been proposed fully leverages subspace and coil information to enable robust reconstruction at high accelerations which leads to shorter scan times. Second, the deep learning prediction is used to initialize an iterative reconstruction instead of being used as the output of the framework, which was demonstrated to guard against potential hallucinations and improve robustness.

Although, this work uses a simple deep learning model (ResUNet^48^) to jump-start the compressed sensing reconstruction, given the flexibility of the Deli-CS framework, various deep learning architecture can be easily integrated to try and improve the quality of the initialization, as well as incorporating B0 and B1 correction strategies. Note that in for rapid whole brain MRF reconstruction. The proposed Deli-CS frameworks aims to fully leverage MRI physics and differs from Gomez et al.^56^ in the following key ways. First, it this work no B1 and B0 correction was included as proposed by Cao et al.^12^ to improve quantification estimates in areas affected by field inhomogeneities. B1 inhomogeneity affects the T2 estimation in the front of the brain, whilst B0 inhomogeneity mostly affects estimates in the lower parts of the brain near the air-tissue interfaces of the sinuses. Future work is to leverage a more advanced calibration scan, such as PhysiCal^57^ which aims to acquire *B*_0_, 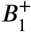, and coil sensitivity information in less than 20 seconds. The 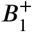 map will enable robust parametric mapping, and the *B*_0_ information can be incorporated into the **A** matrix in equation 2 to alleviate blurring issues related to spiral imaging that is still present even in the highly accelerated spiral trajectory, particularly in regions where the *B*_0_ is large^12^. This is a good fit for the Deli-CS framework, as the network can potentially learn to predict a *B*_0_ corrected image using the non-*B*_0_ corrected input or multiple frequency shifted inputs, which could result in fewer refinement iterations (that would use the full forward model **A** with incorporated *B*_0_). This is beneficial as **A** augmented with *B*_0_ is even more computationally challenging as discussed by Cao et al.^12^, making the traditional LLR reconstruction even harder to perform using realistic time and hardware constraints.

Finally, it would be ideal to port Deli-CS to a more efficient compiled language, e.g. C, which is expected to provide at least another 2 − 3*x* in speed improvement, moving the application towards near real-time reconstruction.

## DATA AVAILABILITY STATEMENT

The code and data used to generate the above results can be found in these repositories and public records:

https://github.com/SetsompopLab/deli-cs

https://github.com/SetsompopLab/MRF

https://zenodo.org/record/7734431

https://zenodo.org/record/7703200

https://zenodo.org/record/7697373

The first repository link provides instructions on how to download and use the data and the second repository.

## ACKNOWLEDGEMENTS

The authors would like to thank Mark Nishimura (ORCID: 0000-0003-3976-254X) for reviewing the code and data shared as part of this article.

## REFERENCES

1. Ma Dan, Gulani Vikas, Seiberlich Nicole, et al. Magnetic Resonance Fingerprinting. Nature. 2013;495(7440):187–192.

2. Kecskemeti Steven, Samsonov Alexey, Hurley Samuel A, Dean Douglas C, Field Aaron, Alexander Andrew L. MPnRAGE: A Technique to Simultaneously Acquire Hundreds of Differently Contrasted MPRAGE Images with Applications to Quantitative T1 Mapping. Magnetic Resonance in Medicine. 2016;75(3):1040–1053.

3. Cao Peng, Zhu Xucheng, Tang Shuyu, Leynes Andrew, Jakary Angela, Larson Peder EZ. Shuffled Magnetization-prepared Multicontrast Rapid Gradient-echo Imaging. Magnetic Resonance in Medicine. 2018;79(1):62–70.

4. Liang Zhi-Pei. Spatiotemporal imaging with partially separable functions. In: 2007 4th IEEE International Symposium on Biomedical Imaging: From Nano to Macro - Proceedings:988–991; 2007.

5. Huang Chuan, Bilgin Ali, Barr Tomoe, Altbach Maria I. T2 Relaxometry with Indirect Echo Compensation from Highly Undersampled Data. Magnetic Resonance in Medicine. 2013;70(4):1026–1037.

6. Velikina Julia V, Alexander Andrew L, Samsonov Alexey. Accelerating MR Parameter Mapping using Sparsity-promoting Regularization in Parametric Dimension. Magnetic Resonance in Medicine. 2013;70(5):1263–1273.

7. Ben-Eliezer Noam, Sodickson Daniel K, Block Kai Tobias. Rapid and Accurate T2 Mapping from Multi–spin-echo Data using Bloch-simulation-based Reconstruction. Magnetic Resonance in Medicine. 2015;73(2):809–817.

8. Zhao Bo, Lu Wenmiao, Hitchens T Kevin, Lam Fan, Ho Chien, Liang Zhi-Pei. Accelerated MR Parameter Mapping with Low-rank and Sparsity Constraints. Magnetic Resonance in Medicine. 2015;74(2):489–498.

9. Velikina Julia V, Samsonov Alexey A. Reconstruction of Dynamic Image Series from Undersampled MRI Data using Data-driven Model Consistency Condition (MOCCO). Magnetic Resonance in Medicine. 2015;74(5):1279–1290.

10. Tamir Jonathan I, Uecker Martin, Chen Weitian, et al. T2-Shuffling: Sharp, multicontrast, volumetric fast spin-echo imaging. Magnetic Resonance in Medicine. 2017;77(1):180–195.

11. Wang Fuyixue, Dong Zijing, Reese Timothy G, et al. Echo Planar Time-Resolved Imaging (EPTI). Magnetic Resonance in Medicine. 2019;81(6):3599–3615.

12. Cao Xiaozhi, Liao Congyu, Iyer Siddharth Srinivasan, et al. Optimized multi-axis spiral projection MR fingerprinting with subspace reconstruction for rapid whole-brain high-isotropic-resolution quantitative imaging. Magnetic Resonance in Medicine. 2022;88(1):133–150.

13. Uecker Martin, Ong Frank, Tamir Jonathan I, et al. Berkeley Advanced Reconstruction Toolbox. Proceedings of the International Society of Magnetic Resonance in Medicine, Toronto, Canada. 2015;23(2486).

14. Beck Amir, Teboulle Marc. A fast iterative shrinkage-thresholding algorithm for linear inverse problems. SIAM Journal on Imaging Sciences. 2009;2(1):183–202.

15. Wajer Ftaw, Pruessmann KP. Major speedup of reconstruction for sensitivity encoding with arbitrary trajectories. Proceedings of the International Society of Magnetic Resonance in Medicine, Toronto, Canada. 2001;.

16. Fessler Jeffrey A, Lee Sangwoo, Olafsson Valur T, Shi Hugo R, Noll Douglas C. Toeplitz-based iterative image reconstruction for MRI with correction for magnetic field inhomogeneity. IEEE Transactions on Signal Processing. 2005;53(9):3393–3402.

17. Baron Corey A, Dwork Nicholas, Pauly John M, Nishimura Dwight G. Rapid compressed sensing reconstruction of 3D non-Cartesian MRI. Magnetic Resonance in Medicine. 2018;79(5):2685–2692.

18. Aggarwal Hemant K, Mani Merry P, Jacob Mathews. MoDL: Model-based deep learning architecture for inverse problems. IEEE transactions on medical imaging. 2018;38(2):394–405.

19. Hammernik Kerstin, Klatzer Teresa, Kobler Erich, et al. Learning a variational network for reconstruction of accelerated MRI data. Magnetic resonance in medicine. 2018;79(6):3055–3071.

20. Küstner Thomas, Fuin Niccolo, Hammernik Kerstin, et al. CINENet: deep learning-based 3D cardiac CINE MRI reconstruction with multi-coil complex-valued 4D spatio-temporal convolutions. Scientific reports. 2020;10(1):1–13.

21. Sandino Christopher M, Lai Peng, Vasanawala Shreyas S, Cheng Joseph Y. Accelerating cardiac cine MRI using a deep learningbased ESPIRiT reconstruction. Magnetic Resonance in Medicine. 2021;85(1):152–167.

22. Wang Xiaoqing, Tan Zhengguo, Scholand Nick, Roeloffs Volkert, Uecker Martin. Physics-based reconstruction methods for magnetic resonance imaging. Philosophical Transactions of the Royal Society. 2021;379(2200):20200196.

23. Hammernik Kerstin, Küstner Thomas, Yaman Burhaneddin, et al. Physics-Driven Deep Learning for Computational Magnetic Resonance Imaging. arXiv preprint arXiv:2203.12215. 2022;.

24. Block Kai Tobias, Uecker Martin, Frahm Jens. Undersampled Radial MRI with Multiple Coils. Iterative Image Reconstruction Using a Total Variation Constraint. Magnetic Resonance in Medicine. 2007;57(6):1086–1098.

25. Lustig Michael, Donoho David, Pauly John M. Sparse MRI: The application of compressed sensing for rapid MR imaging. Magnetic Resonance in Medicine. 2007;58(6):1182–1195.

26. Lustig Michael, Donoho David L., Santos Juan M., Pauly John M. Compressed Sensing MRI. IEEE Signal Processing Magazine. 2008;25(2):72–82.

27. Liu Bo, King Kevin, Steckner Michael, Xie Jun, Sheng Jinhua, Ying Leslie. Regularized Sensitivity Encoding (SENSE) Reconstruction Using Bregman Iterations. Magnetic Resonance in Medicine. 2009;61(1):145–152.

28. Mani Merry, Jacob Mathews, Magnotta Vincent, Zhong Jianhui. Fast iterative algorithm for the reconstruction of multishot non-cartesian diffusion data. Magnetic Resonance in Medicine. 2015;74(4):1086–1094.

29. Fessler Jeffrey A. Optimization Methods for Magnetic Resonance Image Reconstruction: Key Models and Optimization Algorithms. IEEE Signal Processing Magazine. 2020;37(1):33–40.

30. He Kaiming, Zhang Xiangyu, Ren Shaoqing, Sun Jian. Deep residual learning for image recognition. In: :770–778; 2016.

31. Liao Congyu Liao, Cao Xiaozhi, Iyer Siddharth, et al. Mesoscale myelin-water fraction and T1/T2/P D mapping through optimized 3D ViSTa-MRF and stochastic reconstruction with preconditioning. Proceedings of the International Society of Magnetic Resonance in Medicine. 2022;.

32. Desai Arjun D, Gunel Beliz, Ozturkler Batu M, et al. VORTEX: Physics-Driven Data Augmentations for Consistency Training for Robust Accelerated MRI Reconstruction. arXiv preprint arXiv:2111.02549. 2021;.

33. Desai Arjun D, Ozturkler Batu M, Sandino Christopher M, et al. Noise2recon: A semi-supervised framework for joint mri reconstruction and denoising. arXiv preprint arXiv:2110.00075. 2021;.

34. Darestani Mohammad Zalbagi, Heckel Reinhard. Accelerated MRI with un-trained neural networks. IEEE Transactions on Computational Imaging. 2021;7:724–733.

35. Darestani Mohammad Zalbagi, Chaudhari Akshay S, Heckel Reinhard. Measuring robustness in deep learning based compressive sensing. In: :2433–2444PMLR; 021.

36. Darestani Mohammad Zalbagi, Liu Jiayu, Heckel Reinhard. Test-Time Training Can Close the Natural Distribution Shift Performance Gap in Deep Learning Based Compressed Sensing. arXiv preprint arXiv:2204.07204. 2022;.

37. Ong Frank, Lustig Michael. SigPy: A Python Package for High Performance Iterative Reconstruction. Proceedings of the International Society of Magnetic Resonance in Medicine, Montréal, Canada. 2019;4819.

38. Fessler J.A., Sutton B.P.. Nonuniform fast Fourier transforms using min-max interpolation. IEEE Transactions on Signal Processing. 2003;51(2):560–574.

39. Pruessmann Klaas P, Weiger Markus, Scheidegger Markus B, Boesiger Peter. SENSE: sensitivity encoding for fast MRI. Magnetic Resonance in Medicine: An Official Journal of the International Society for Magnetic Resonance in Medicine. 1999;42(5):952–962.

40. Uecker Martin, Lai Peng, Murphy Mark J, et al. ESPIRiT - an eigenvalue approach to autocalibrating parallel MRI: where SENSE meets GRAPPA. Magnetic resonance in medicine. 2014;71(3):990–1001.

41. Norbeck Ola, Sprenger Tim, Avventi Enrico, et al. Optimizing 3D EPI for Rapid T1-Weighted Imaging. Magnetic Resonance in Medicine. 2020;84(3):1441–1455.

42. Kim Daeun, Cauley Stephen F., Nayak Krishna S., Leahy Richard M., Haldar Justin P. Region-optimized virtual (ROVir) coils: Localization and/or suppression of spatial regions using sensor-domain beamforming. Magnetic Resonance in Medicine. 2021;86(1):197–212.

43. Pipe James G., Menon Padmanabhan. Sampling density compensation in MRI: Rationale and an iterative numerical solution. Magnetic Resonance in Medicine. 1999;41(1):179–186.

44. Cruz Gastao, Jaubert Olivier, Schneider Torben, Botnar Rene M, Prieto Claudia. Rigid Motion-Corrected Magnetic Resonance Fingerprinting. Magnetic Resonance in Medicine. 2019;81(2):947–961.

45. Cruz Gastão, Jaubert Olivier, Qi Haikun, et al. 3D Free-Breathing Cardiac Magnetic Resonance Fingerprinting. Nuclear Magnetic Resonance in Biomedicine. 2020;33(10):e4370.

46. Bradski G.. The OpenCV Library. Dr. Dobb’s Journal of Software Tools. 2000;.

47. Ying Leslie, Sheng Jinhua. Joint Image Reconstruction and Sensitivity Estimation in SENSE (JSENSE). Magnetic Resonance in Medicine. 2007;57(6):1196–1202.

48. Zhang Zhengxin, Liu Qingjie, Wang Yunhong. Road Extraction by Deep Residual U-Net. IEEE Geoscience and Remote Sensing Letters. 2018;15(5):749–753.

49. Paszke Adam, Gross Sam, Massa Francisco, et al. PyTorch: An Imperative Style, High-Performance Deep Learning Library. In: Wallach H., Larochelle H., Beygelzimer A., Alché-Buc F., Fox E., Garnett R., eds. Advances in Neural Information Processing Systems 32, Curran Associates, Inc. 2019 (pp. 8024–8035).

50. Falcon William. PyTorch Lightning. GitHub. Note: https://github.com/PyTorchLightning/pytorch-lightning. 2019;3.

51. Kingma Diederik P., Ba Jimmy. Adam: A Method for Stochastic Optimization. 2017.

52. Zhang Y., Brady M., Smith S.. Segmentation of brain MR images through a hidden Markov random field model and the expectation-maximization algorithm. IEEE Transactions on Medical Imaging. 2001;20(1):45–57.

53. Smith Stephen M. Fast robust automated brain extraction. Human Brain Mapping. 2002;17(3):143–155.

54. Demir Serdest, Clifford Bryan, Lo Wei-Ching, et al. Optimization of magnetization transfer contrast for EPI FLAIR brain imaging. Magnetic Resonance in Medicine. 2022;87(5):2380–2387.

55. Kim Daeun, Wisnowski Jessica L., Nguyen Christopher T., Haldar Justin P. Multidimensional correlation spectroscopic imaging of exponential decays: From theoretical principles to in vivo human applications. NMR in Biomedicine. 2020;33(12):e4244. e4244 nbm.4244.

56. Gómez Pedro A, Cencini Matteo, Golbabaee Mohammad, et al. Rapid three-dimensional multiparametric MRI with quantitative transient-state imaging. Scientific reports. 2020;10(1):1–17.

57. Iyer Siddharth, Liao Congyu, Li Qing, et al. PhysiCal: A rapid calibration scan for B0, B1+, coil sensitivity and Eddy current mapping. In: :661; 2020.

